# Alternative splicing of pericentrin contributes to cell cycle control in cardiomyocytes

**DOI:** 10.1101/2021.04.19.440474

**Authors:** Jakob Steinfeldt, Robert Becker, Silvia Vergarajauregui, Felix B. Engel

## Abstract

Induction of cardiomyocyte proliferation is a promising option to regenerate the heart. Thus, it is important to elucidate mechanisms that contribute to the cell cycle arrest of mammalian cardiomyocytes. Here, we assessed the contribution of the pericentrin (Pcnt) S isoform to the cell cycle arrest in postnatal cardiomyocytes. Immunofluorescence staining of Pcnt isoforms combined with siRNA-mediated depletion indicates that Pcnt S preferentially localizes to the nuclear envelope, while the Pcnt B isoform is enriched at centrosomes. This is further supported by the localization of ectopically expressed FLAG-tagged Pcnt S and Pcnt B in postnatal cardiomyocytes. Analysis of centriole configuration upon Pcnt depletion revealed that Pcnt B but not Pcnt S is required for centriole cohesion. Importantly, ectopic expression of Pcnt S induced centriole splitting in a heterologous system, ARPE-19 cells, and was sufficient to impair DNA synthesis in C2C12 myoblasts. Moreover, Pcnt S depletion enhanced serum-induced cell cycle re-entry in postnatal cardiomyocytes. Analysis of mitosis, binucleation rate, and cell number suggests that Pcnt S depletion promotes progression of postnatal cardiomyocytes through the cell cycle resulting in cell division. Collectively, our data indicate that alternative splicing of Pcnt contributes to the establishment of cardiomyocyte cell cycle arrest shortly after birth.

## 1. Introduction

Ischemic heart disease (IHD) is the leading cause of death worldwide [1]. Currently IHD is treated by reopening occluded vessels (acute percutaneous coronary intervention) to reestablish perfusion and pharmacologically (e.g. beta blockers) to minimize cardiac remodeling and further deterioration of heart function. Except heart transplantation, which is limited due to a shortage in organ donors, there are no therapies available to effectively reverse heart injury. One promising approach to regenerate lost heart muscle tissue is the modulation of developmental pathways to induce endogenous proliferation of differentiated cardiomyocytes [2].

Mammalian cardiomyocytes stop to proliferate shortly after birth. The cells progress through a last cell cycle, which results in polyploidization or binucleation and subsequently cell cycle arrest [3–5]. Whether this cell cycle arrest is permanent or can be reversed was long debated. In 2005, it has been demonstrated that adult mammalian cardiomyocytes can be induced to undergo cell division [6]. Subsequently, a large number of studies has been published supporting the idea that cardiac regeneration can be achieved by induction of cardiomyocyte proliferation [7,8]. Yet, it has also been suggested that induction of postnatal cardiomyocyte cell division is associated with improper distribution of microtubules and might result in chromosome mis-segregation due to the formation of pseudo-bipolar spindles [9,10]. Overall, there are issues regarding technologies used to prove cardiomyocyte proliferation, the efficiency of cardiomyocyte proliferation induction, and the causality between cardiomyocyte proliferation and improved cardiac function [7,8,11].

Pericentrin (Pcnt) is a multifunctional scaffold protein that binds to a large variety of centrosomal proteins [12,13]. Consequently, Pcnt regulates a large number of centrosomal functions such as control of cell cycle progression, mitotic spindle organization and orientation, oriented cell division, and cell fate determination [13,14]. There are two major alternative splice isoforms of Pcnt, Pcnt B (also known as kendrin) and the shorter Pcnt S, which lacks part of the N-terminal region of Pcnt B [15]. Pcnt B is localized at the centrosome in the vast majority of cell types throughout development. In contrast, Pcnt S is expressed predominantly in the adult heart, skeletal muscle, and testis [15,16]. Upregulation of Pcnt S is in cardiomyocytes associated with loss of centrosome integrity and cell cycle arrest [17]. After birth, various centrosome proteins, such as Pcnt S, are localized to the nuclear envelope and the paired centrioles, the center of the centrosome, lose cohesion. This process results in split centrioles (> 2 μm apart from each other) and the establishment of a non-centrosomal microtubule organizing center (MTOC) at the nuclear envelope [17–19]. Similarly to cardiomyocytes, skeletal myoblasts form a nuclear envelope MTOC during differentiation [17,18]. Besides three studies describing the existence of Pcnt S in the adult heart and skeletal muscle and the localization of Pcnt S at the nuclear envelope in cardiomyocytes [15–17], nothing is known about Pcnt S. Considering the role of Pcnt B in cell cycle control and the association of Pcnt S with differentiation and cell cycle arrest in striated muscle cells, we hypothesized that alternative splicing of Pcnt contributes to the establishment of the cell cycle arrest of mammalian cardiomyocytes. The importance of Pcnt for proper human development is underlined by a number of human disorders linked to Pcnt mutations [13,20].

Here, we present evidence that alternative splicing of Pcnt contributes to cell cycle control in mammalian cardiomyocytes. We show by antibody staining in combination with siRNA-mediated Pcnt depletion and ectopic expression of tagged Pcnt isoforms that Pcnt B preferentially localizes at the centrosome and Pcnt S at the nuclear envelope. While ectopic expression of Pcnt S induces centriole splitting and inhibits cell cycle progression in skeletal myoblasts, siRNA-mediated depletion of Pcnt S promotes serum-induced cell cycle re-entry, progression through mitosis and cell division of cardiomyocytes.

## 2. Materials and Methods

### 2.1. Isolation and cell culture of postnatal cardiomyocytes

Mammalian ventricular cardiomyocytes were isolated on day 3 after birth (P3) as previously described [6], seeded on 1 mg/ml fibronectin (Sigma)-coated glass coverslips and cultured in cardiomyocyte medium (DMEM-F12, Glutamax TM-I, 3 mM Na-pyruvate, 0.2% bovine serum albumin (BSA), 0.1 mM ascorbic acid, 0.5% Insulin-Transferrin-Selenium (100x, Life Technologies), and penicillin/streptomycin (100 U/mg/ml)) containing horse serum (HS, 5% for testing depletion efficiency and effect on centriole cohesion, 1% for assessing the effect on cell cycle progression) for 24 h prior to experimentation.

### 2.2. Cell culture of C2C12 myoblasts and ARPE-19 adult retinal pigment epithelial cells

Cells were maintained in a humidified atmosphere containing 5% CO2 at 37°C and subcultured every two to three days. Culture medium consisted of DMEM/F12 supplemented with GlutaMAX, 4.5 mg/ml D-glucose (high glucose) (Thermo Fisher/Gibco #31331-093), 10% fetal bovine serum (FBS, Biowest) and 50 mg/ml gentamycin (Serva). For transfection, cells were seeded on 12 mm glass coverslips in a 24-well plate at a density of 10.000 (C2C12) or 50.000 cells (ARPE-19)/well. For centriole analysis, cells were fixed 24 h after transfection.

### 2.3. siRNA knockdown

Lipofectamine RNAiMAX reagent (ThermoFisherScientific, Waltham, Massachusetts, US) was utilized to transfect cardiomyocytes 24 hours post-seeding with siPcntS (50 nM or 100 nM as indicated, 5’-CAUAUGUUCUUGUAUAAAAtt-3’), an siRNA targeting the Pcnt S-specific 5’UTR or with siPcntTotal (200 nM, 5’-CAGGAACUCACCAGAGAC-GAA-3’), an siRNA targeting the C-terminal part of Pcnt, which is common to Pcnt B and Pcnt S. After transfection, cells were incubated for 72 h in medium containing 5% HS for testing depletion efficiency and effects on centriole cohesion. 10% FBS was utilized for assessing the effect on cell cycle progression. siRNA-mediated depletion efficiency was assessed by determining the median intensity of total Pcnt at the nucleus in P3 cardiomyocytes. Pcnt signal from the centrosome was considered noise. The baseline was set as the median nuclear Pcnt signal of non-myocytes – which contains overlapping centriolar signal as well- and subtracted from all measurements. To determine the effect of siPcntS on Pcnt B expression, the maximum intensity of the Pcnt B signal per cardiomyocyte was measured as an approximation of the centrosomal Pcnt B signal.

### 2.4. Analysis of DNA incorporation

To detect entry into S phase, 30 μM EdU was added to the medium post-transfection (cardiomyocytes: 48 h, C2C12 cells: 45 h). After 3 h or 24 h C2C12 cells and cardiomyocytes were fixed, respectively. EdU incorporation was determined utilizing the kit Click-iT^®^ EdU Alexa Fluor^®^ 488 (Life Technologies) according to the manufacturer’s protocol.

### 2.5. Plasmids

p3xFLAG-CMV10-eGFP-hPCNTB (17027 bp, FLAG-Pcnt B) was a gift from Kunsoo Rhee [21]. This construct was used to create a construct to ectopically express Pcnt S by amplifying the backbone and the C-terminal part of Pcnt B (#1 Backbone FV: gccacccgatgattaaacagGATATCGAGCAGAAACTCATCTCTG, #2 Backbone RV: agcggccgcCTTGTACAGCTCGTCCATGCCGAGAG, #3 Pcnt S FV: ATGCTCAAGGCCGACGTCAACCTGT, #4 Pcnt S RV: atgagtttctgctcgatatcCTGTTTAATCATCGGGTGGCAGGAT, #5 Pcnt B 5’ specific FV: agctgtacaagGCGGCCGCT, #6 Pcnt B 5’ specific RV: TTGACGTCGGCCTTGAGCAT). Linearized vectors were fused using the Cold Fusion Cloning Kit and transformed in E. coli to obtain the construct p3xFLAG-CMV10-eGFP-hPCNTS (13601bp, FLAG-Pcnt S) which was controlled by sequencing. To obtain the plasmids PcntS-T2A-eGFP and PcntB-T2A-eGFP, the coding sequences of human Pcnt B and Pcnt S were subcloned into peGFP-N1 (Takara Clontech) using the NEBuilder HiFi DNA Assembly Master Mix (New England Biolabs) according to manufacturer’s instructions. Fragments for the assembly reaction were generated with CloneAmp™ HiFi PCR Premix (Takara Clontech) using the following primers: #1 Pcnt B part 1 forward: gggatccaccggtcgccaccATGGAAGTTGAGCAAGAGCAG; #2 Pcnt B part 1 reverse: ctgcagactgCCAGCCTGACTGTCGCTG; #3 Pcnt B part 2 forward: gtcaggctggCAGTCTGCAGAGCGAGCTG; #5 Pcnt S forward: gggatccaccggtcgccaccATGCTCAAGGCCGACGTC; #6 Pcnt B part 2 + Pcnt S reverse: cgtcaccgcatgttagcagacttcctctgccctctccactgccCTGTTTAATCATCGGGTGGC; #7 peGFP-N1 forward: aagtctgctaacatgcggtgacgtcgaggagaatcctggcccaATGGTGAGCAAGGGCGAG; #8 peGFP-N1 reverse: GGTGGCGACCGGTGGATC. Primers #6 and #7 contain the coding sequence for a T2A site, enabling expression of the Pcnt isoforms and eGFP as separate proteins. Correct assembly was confirmed by restriction digest as well as sequencing.

### 2.6. Plasmid transfection

Cells were transfected with plasmids utilizing Lipofectamine LTX and LTX Plus Reagent (Thermo Fisher; 500 ng plasmid DNA, 1 μl (cardiomyocytes, ARPE19) or 2 μl (C2C12) LTX reagent and 1 μl (cardiomyocytes) or 0.5 μl (C2C12, ARPE19) LTX Plus reagent, per well of a 24-well plate). DNA-liposome complexes were assembled in Opti-MEM (Thermo Fisher) for 20 min and then applied to the culture medium. Cardiomyocyte were analyzed 48 h post-transfection.

### 2.7. Immunofluorescence and microscopy

Fixation using pre-chilled methanol for 3 min at −20°C was used for testing the depletion efficiency and effects on centriole cohesion in cardiomyocytes. 4% formaldehyde (Carl Roth, Karlsruhe, Germany) for 10 min at room temperature was used for assessing the effects on cell cycle. 10% FBS in PBS + 0.1% saponin + 0.03% sodium azide was used as blocking buffer (20 min at 20°C). Formalin-fixed cells were permeabilized prior to antibody staining with 0.5% Triton X-100 (Sigma)/PBS for 10 min at room temperature. Primary antibodies: rabbit anti-Pcnt (1:500; MmPeriC1) against both B and S isoforms was produced as previously described [22], mouse anti-Pcnt against Pcnt B (1:500; MmPeri N-term clone 8D12) was made against the first 233 amino acids of mouse PCNT B (AN: NP_032813 or BAF36559), mouse anti-PCM1 (1:500, sc-398365, Santa Cruz Biotechnology, Dallas, TX, USA), goat anti-troponin I (1:250, ab56357, Abcam, Cambridge, UK), mouse anti-コ-tubulin (1:500, sc-51715, Santa Cruz Biotechnology, Dallas, TX, USA), anti-Ki67 (1:250, ab8191, Abcam, Cambridge, UK). Secondary antibodies: ALEXA 488-, ALEXA 594-, and ALEXA 647-conjugated antibodies (1:500, Life Technologies, Carlsbad, CA, USA). DNA was stained with 0.5 μg/ml DAPI (4′,6′-diamidino-2-phenylindole) (Sigma). Images were captured on a Zeiss LSM 800 Confocal Fluorescence Microscope (Zeiss, Oberkochen, Germany), using 10x, 20x or 63x objectives and the ZEISS Blue software.

### 2.8. Image preparation and analysis

Images were arranged with Fiji [23], a distribution of ImageJ (Public Domain), custom Python scripts (Python Software Foundation, https://www.python.org/), the R package ggplot2 [24], and Adobe Illustrator (Adobe, San Jose, CA, USA). For image analysis Fiji was used for preprocessing, custom Cellprofiler [25] pipelines for the extraction of features and measurements in a SQLite database and the Cellprofiler Analyst [26] for manually labeling the training dataset. Random Forest classifier were trained in Python on the training set and performance was evaluated via 5-fold cross-validation. The classifier was utilized to classify all cells automatically for the proceeding analysis and the classification results were written in the SQLite database. Centrioles were identified by visualizing the log normalized intensity values while classifying the images.

### 2.9. Statistical analysis

Image intensities were normalized per experiment. Binucleated cardiomyocytes were excluded from the quantification of centriole cohesion, Ki67 ratio, EdU ratio and cardiomyocyte count, to control for false positive proliferation. H0 was defined as the group distributions being equal, H1 was defined as them being different. Statistical analysis was performed using R 3.6.1 [27] and reading the measurements and classifications from an SQLite database. Data of at least three biological independent experiments with at least 10 technical replicates each was either expressed as a bar plot with mean ± SD or Box-Whiskers-Plot. To test for differences in the mean between groups and to take into account the biological and technical replications, we used a nested ANOVA design by fitting a linear mixed effects models with restricted maximum likelihood using the R package nlme [28]. Bonferroni-corrected p values were obtained by comparing the linear hypothesis of the mixed effects model. p values were reported as significant, when they were lower than the alpha level of 0.05. Additional bayesian statistical analysis was performed by using the BayesFactor package [29]. To compute the posterior distribution of the group wise treatment effect we used Marcov-Chain-Monte-Carlo sampling and a noninformative Jeffreys prior for the variance and a standard Cauchy prior for the mean.

## 3. Results

### 3.1. Pcnt B preferentially localizes at the centrosome and Pcnt S at the nuclear envelope

Previously, it has been reported that “Pericentrin S, and not Pericentrin B, is the predominant Pericentrin isoform at the nuclear envelope in P3-isolated cardiomyocytes” [17]. This conclusion was drawn based on an antibody specific for Pcnt B and an antibody detecting both, Pcnt B and S. To address specifically the location and function of Pcnt S, we designed a siRNA specific for the 5’ UTR of Pcnt S (siPcntS) and a siRNA targeting the C-terminal part of Pcnt, which is common to Pcnt B and Pcnt S (siPcntB+S) (Figure S1). Knockdown efficiency in ventricular cardiomyocytes isolated from three days old rats (P3 cardiomyocytes) was validated by semi-quantitative immunofluorescence analysis of median intensity of nuclear Pcnt B+S signal (Figure 1A,B) and the Pcnt B signal at the centrosome (Figure 1C,D).

**Figure 1.**
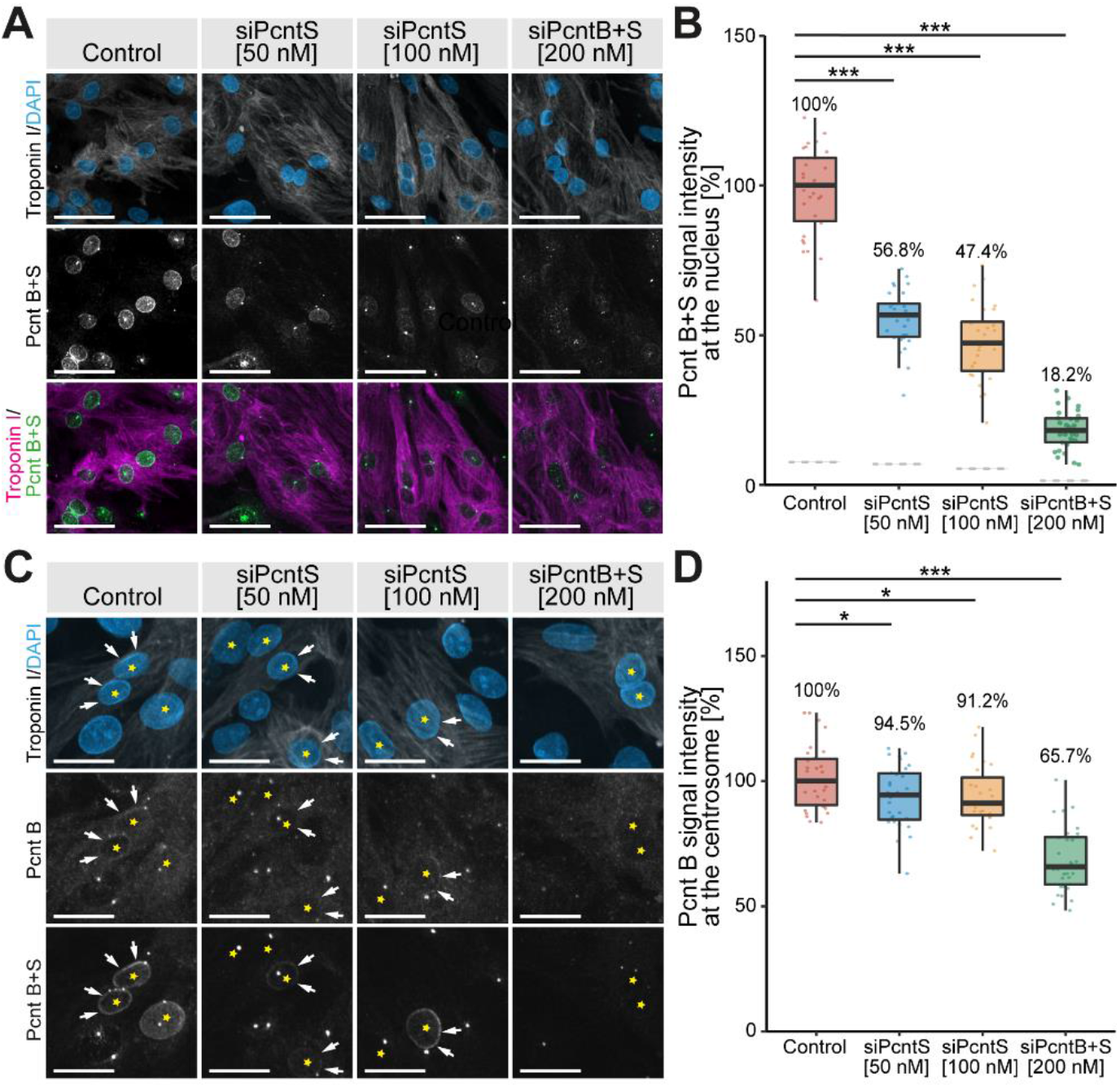
Immunofluorescence analysis of Pcnt S and Pcnt B localization. P3 cardiomyocytes were transfected with siRNAs to deplete Pcnt S or Pcnt S + B as indicated. (**A**) Immunofluorescence analysis of Pcnt expression in cardiomyocytes (troponin I) utilizing an antibody binding to both Pcnt S and Pcnt B (Pcnt B+S). Nuclei were visualized with DAPI (DNA). (**B**) Semi-quantitative analysis of median intensity of nuclear Pcnt B+S signal in a. (**C**) Immunofluorescence analysis of Pcnt expression in cardiomyocytes (troponin I) utilizing an antibody binding to both Pcnt S and Pcnt B (Pcnt B+S) and an antibody detecting specifically Pcnt B. Nuclei were visualized with DAPI (DNA). (**D**) Semi-quantitative analysis of maximum intensity of the Pcnt B signal per cardiomyocyte in **C** as an approximation of the centrosomal Pcnt B signal. Yellow asterisk: cardiomyocyte nucleus. White arrows: nuclear envelope of cardiomyocytes. For the experiments ≥ 2000 cardiomyocytes were analyzed per experimental condition. Scale bars: 50 μm. Data are mean ± SD, n = 3, p < 0.05, **: p < 0.01, ***: p < 0.001.

The nuclear signal of Pcnt contained some noise from overlapping centriolar signals. To overcome this noise, the median Pcnt signal from the nucleus of non-myocytes, which do not have Pcnt at the nuclear envelope, was subtracted from all measurements. In three independent experiments a total of 7325 cardiomyocytes (troponin I-positive) was analyzed. In control experiments, we detected for both, Pcnt B+S as well as Pcnt B, a positive signal at the nuclear envelope. Upon siRNA-mediated knockdown of Pcnt S or Pcnt B+S, a significant reduction of nuclear Pcnt B+S signal was observed in all treatment groups (50 nM siPcntS: 56.8%, 100 nM siPcntS: 47.4%, 200 nM of siPcntB+S: 18.2%, ANOVA: p < 0.001, Figure 1A,B). Notably, depletion of Pcnt S had no marked effect on the weak but constant Pcnt B expression at the nuclear envelope (Figure 1C). These data indicate that Pcnt B is like Pcnt S present at the nuclear envelope.

To determine whether siPcntB+S is sufficient to deplete Pcnt B from the centrosome, we compared the maximum intensity of the Pcnt B signal per cardiomyocyte as an approximation of the centrosomal Pcnt B signal, as Pcnt B at the centrosome represents by far the strongest signal in a cell. siPcntB+S-mediated depletion of both Pcnt isoforms resulted in a clear reduction of the maximum Pcnt intensity to 65.7% (p < 0.001). In contrast, siPcntS-mediated depletion resulted only in a slight reduction to 94.5% (50 nM, p < 0.05) or 91.2% (100 nM, p < 0.05), respectively (Figure 1C,D). Similarly, siPcntS had no obvious effect on the Pcnt B+S signal at the centrosomes (Figure 1C, lowest row). These data suggest that if at all only small amounts of Pcnt S are localized at the centrosome and that Pcnt S depletion does not affect the expression of Pcnt B at the centrosome.

Taken together, the siRNA-related data indicate that siPcntS efficiently depletes Pcnt S and does not interfere with Pcnt B localization. In contrast, siPcntB+S is not efficient in depleting centrosomal Pcnt, even though it resulted in a notable reduction of centrosomal Pcnt.

To validate our initial conclusions that Pcnt S as well as Pcnt B can localize to the centrosome as well as nuclear envelope, FLAG-tagged Pcnt B or Pcnt S were ectopically expressed in P3 cardiomyocytes. Anti-FLAG staining revealed that Pcnt B-FLAG and Pcnt S-FLAG localized to the centrioles/centrosome (γ-tubulin-positive) as well as the nuclear envelope (Figure 2). Notably, the signal for Pcnt S-FLAG was more pronounced at the nuclear envelope than that for Pcnt B-FLAG.

**Figure 2.**
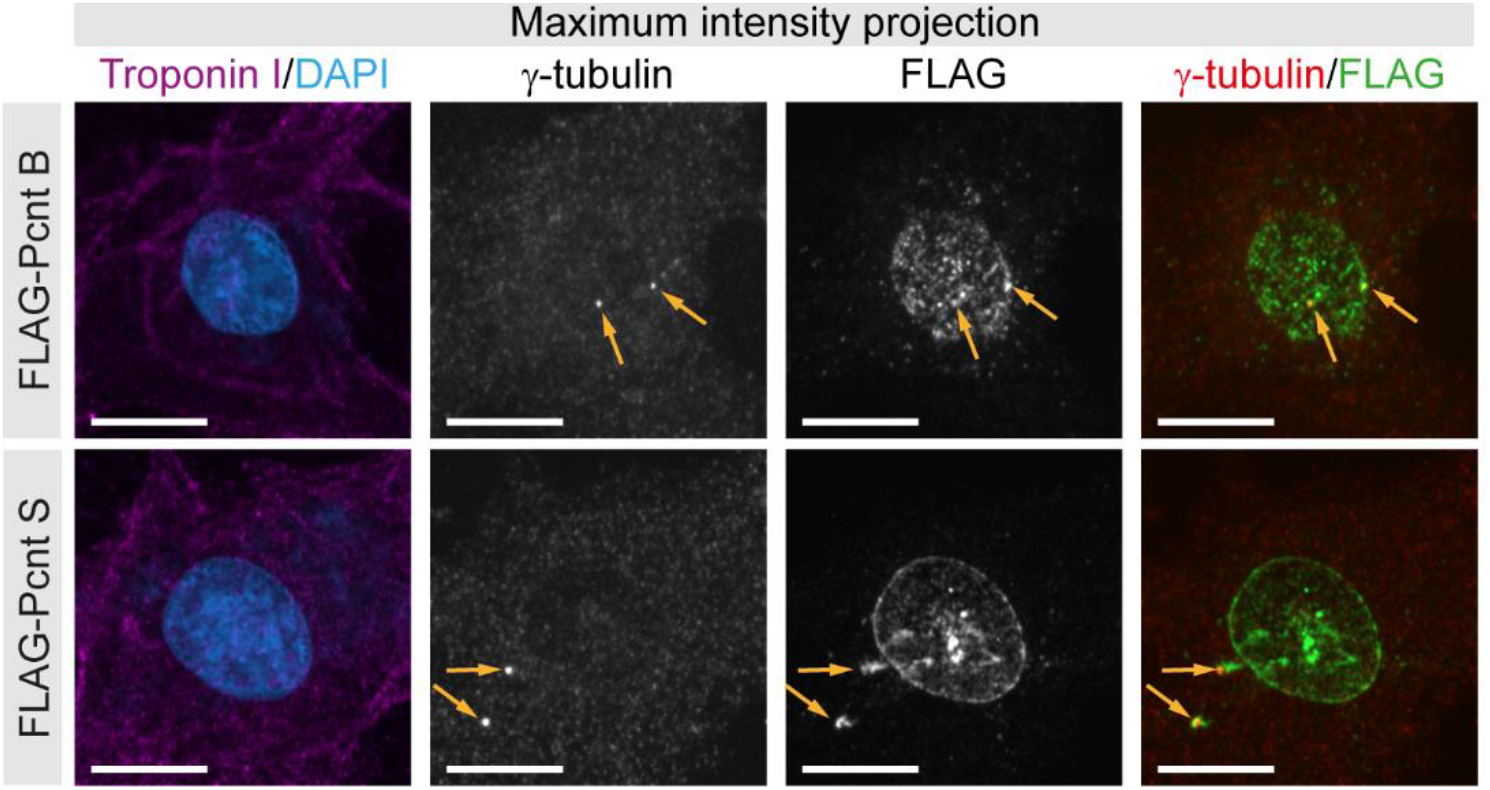
Localization of ectopically expressed Pcnt S and Pcnt B. FLAG-tagged Pcnt B (FLAG-Pcnt B) or Pcnt S (FLAG-Pcnt S) were expressed in P3 cardiomyocytes (troponin I) and their localization was assessed by staining for FLAG as well as centrioles/centrosome (γ-tubulin). Nuclei were visualized with DAPI (DNA). Orange arrows: centrioles/centrosomes. Scale bars: 10 μm.

Collectively, our data indicate that in postnatal cardiomyocytes the dominant Pcnt isoform at the nuclear envelope is Pcnt S and at the centrosome Pcnt B. Yet, our data suggest that both isoforms can localize to the nuclear envelope as well as the centrosome.

### 3.2. Pcnt S is not required for centriole cohesion

It is well known that Pcnt is required for centriole cohesion (i.e. centriole splitting) [30,31]. Yet, the role of Pcnt S in this process is unclear. Notably, loss of centriole cohesion in cardiomyocytes and upregulation of Pcnt S occur at around the same time in development [17]. To determine whether depletion of Pcnt S affects centriole cohesion, P3 cardiomyocytes were transfected with the specific siRNAs, stained for Pcnt B+S, and analyzed for centriole configuration (paired or split) whereby centrioles were considered split if the distance between them exceeded 2 μm (Figure 3A). Since neither siPcntS nor siPcntB+S completely eliminates the Pcnt signal from the centrosome (Figure 1), we utilized Pcnt B+S staining to distinguish between paired and split centrioles (Figure 3A). Analysis of three independent experiments and a total of 3725 cardiomyocytes revealed that Pcnt S depletion had no significant effect on centriole splitting compared to siRNA control (paired centrioles: control: 10.5%, 50 nM siPcntS: 12.1%, 100 nM siPcntS: 10.9%, Figure 3B,C). In contrast, depletion of both isoforms significantly decreased the number of cardiomyocytes with paired centrioles (200 nM siPcntTotal: 5.0%, p < 0.01, Figure 3B,C). These data confirm that Pcnt B is required for centriole cohesion and suggest that Pcnt S is not required for centriole cohesion.

**Figure 3.**
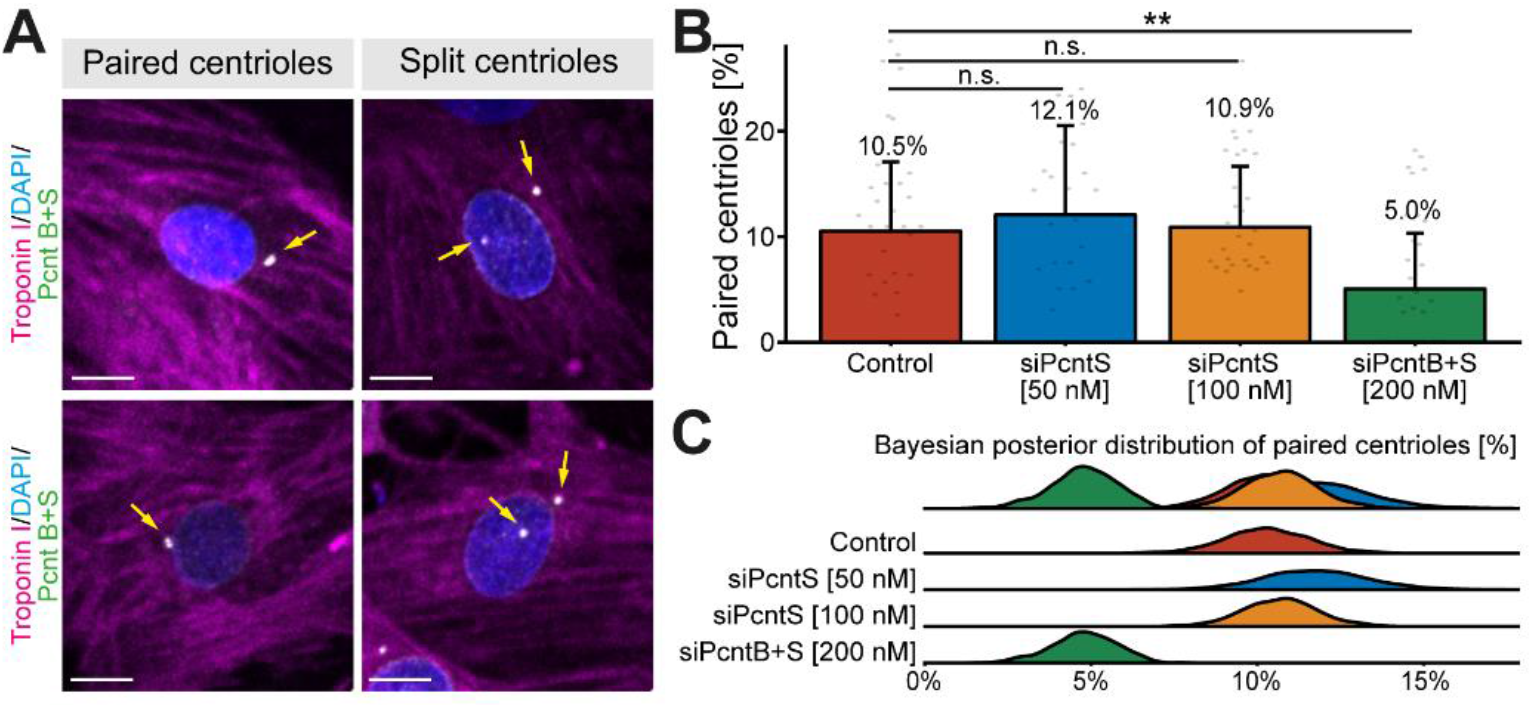
Effect of Pcnt depletion on centriole configuration. (**A**) Representative examples of P3 cardiomyocytes (troponin I) with paired and split centrioles detected by immunofluorescence analysis of Pcnt S and Pcnt B (Pcnt B+S). Nuclei were visualized with DAPI (DNA). (**B**) Quantitative analysis of cardiomyocytes with paired centrioles for control cells and upon Pcnt isoform depletion as indicated. (**C**) Bayesian posterior distribution for the results in b. Scale bars: 10 μm. Yellow arrows: centrioles/centrosomes. Data are mean ± SD, n = 3, n.s.: p > 0.05, **: p < 0.01.

### 3.3. Ectopic expression of Pcnt S induces centriole splitting and centriolar γ-tubulin reduction

To determine whether Pcnt S is sufficient to induce centriole splitting, we expressed Pcnt S-T2A-eGFP or Pcnt B-T2A-eGFP and analyzed centriole configuration in adult retinal pigment epithelial cells (ARPE-19), which do not exhibit centriole splitting (Figure 4A). The introduction of the T2A sequence should result in the expression of the Pcnt isoforms and GFP as separate proteins. Consequently, GFP localized in Pcnt B-T2A-eGFP-transfected cells in the cytoplasm (Figure S2). Yet, in PCNT S-T2A-eGFP-transfected cells GFP accumulated in a centrosome-like manner indicating that the T2A sequence is not functional in PCNT S-T2A-eGFP (sequence has been verified by sequencing) (Figure S2). Immunostaining of Pcnt in Pcnt B-T2A-eGFP- and PCNT S-T2A-eGFP-transfected cells demonstrated that in both cases Pcnt is overexpressed (Figure S2). In order to determine whether any of the two Pcnt isoforms induces centriole splitting, centriolar γ-tubulin signal was analyzed (Figure 4A). As previously introduced, cells were classified by their centriole status and centrioles were considered split if the distance between them exceeded 2 μm. Analysis of 2469 cells from three independent experiments showed increased centriole splitting after overexpression of both Pcnt isoforms (Figure 4B,C). While overexpression of Pcnt B-T2A-eGFP increased centriole splitting in the GFP^+^ over the GFP-cells from 6.3% to 10.8% (p = 0.055), the difference was much more pronounced after ectopic expression of Pcnt S-T2A-eGFP with 5.0% to 15.8% (p > 0.001). Thus, ectopic expression of Pcnt S-T2A-eGFP induced ~46% more centriole splitting than overexpression of Pcnt B-T2A-eGFP.

**Figure 4.**
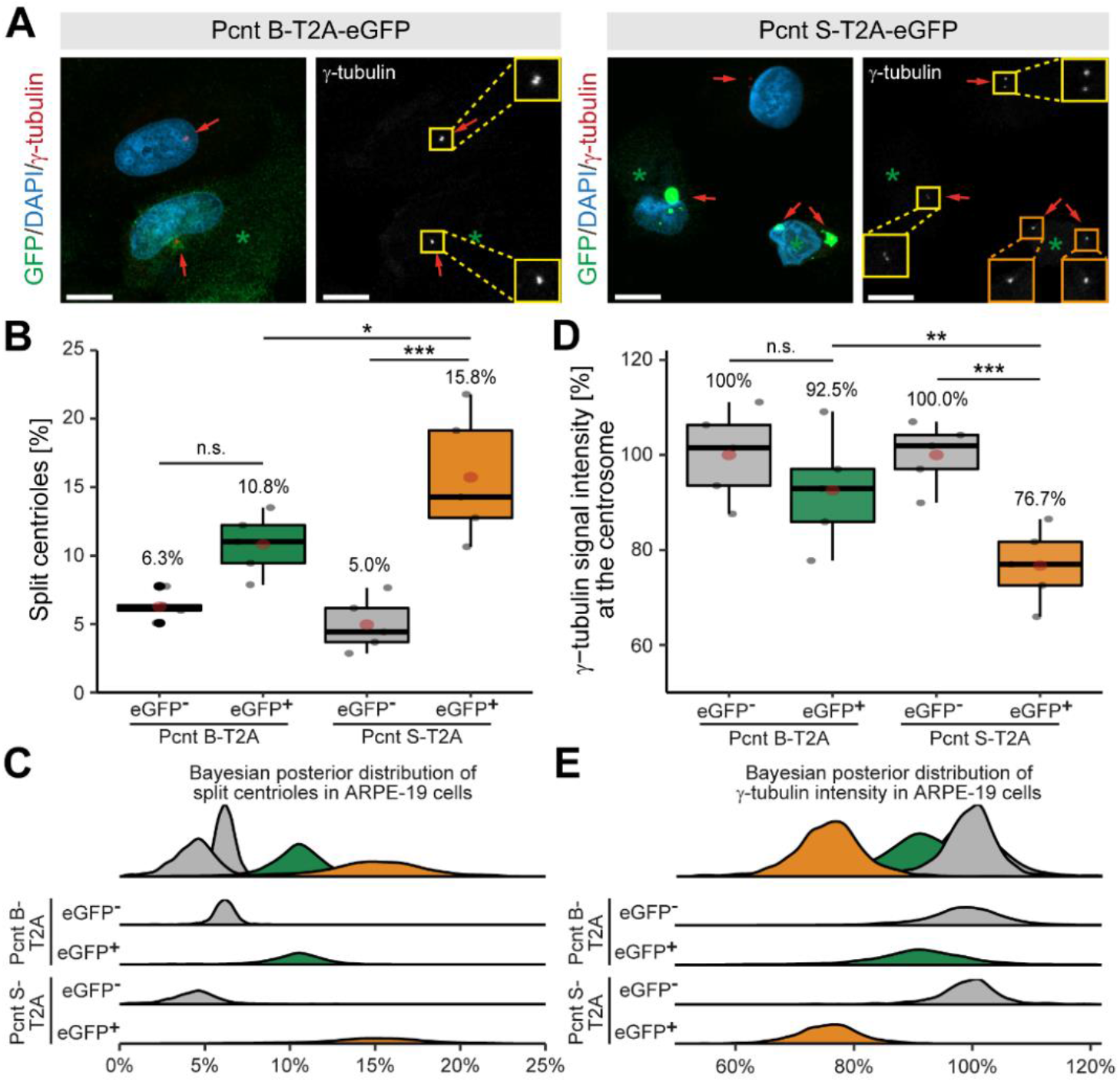
Ectopic expression of Pcnt S induces centriole splitting and reduction of centriolar γ-tubulin in ARPE-19 adult retinal pigment epithelial cells. (**A**) Representative examples of ARPE-19 cells transfected with the bi-cistronic plasmid encoding Pcnt B-T2A-eGFP or Pcnt S-T2A-eGFP (GFP, green) and immunostained with anti-γ-tubulin antibody (centrioles, red). Nuclei were visualized with DAPI (DNA, blue). Yellow box: paired centrioles. Orange box: split centrioles. (**B**,**D**) Quantitative analysis of the percentage of cells showing centriole splitting (**B**) and normalized γ-tubulin intensity at the centrioles compared to non-transfected cells (GFP-) (**D**) upon Pcnt isoform expression as indicated. (**C**,**E**) Bayesian posterior distribution for the data in **B** and **D**. Green asterisk: GFP+ cell. Red arrow: centriole: Scale bars: 10 μm. Data are median ± quartiles and mean (red point), n = 3, n.s.: p > 0.05, *: p < 0.05, ***: p < 0.001.

While examining centriole splitting, we noticed that γ-tubulin signal appeared to be reduced in Pcnt S-T2A-eGFP-transfected ARPE19 cells (Figure 4A). Semi-quantitative analysis confirmed that centriolar γ-tubulin signal intensity in GFP+ cells was notably lower compared to the GFP-cells (100% to 76.7%, p < 0.001) in cultures transfected with Pcnt S-T2A-eGFP (Figure 4D,E). In contrast, γ-tubulin intensity was only mildly reduced in GFP+ cells after overexpressing Pcnt B-T2A-eGFP from 100% to 92.5% (p = 0.73). This indicates that ectopic Pcnt S expression interferes with γ-tubulin localization to centrioles/centrosomes. Taken together, our data suggest that ectopic expression of Pcnt S can induce centriole splitting in ARPE-19 cells. This further indicates that the developmental upregulation of Pcnt S might contribute to the postnatal cell cycle arrest in mammalian cardiomyocytes.

### 3.4. Ectopic Pcnt S expression impairs DNA synthesis

Considering that Pcnt S is upregulated when cardiomyocyte cell cycle progression is arrested, we wondered whether Pcnt S overexpression inhibits cell cycle progression. As cardiomyocyte proliferation cannot be maintained in fetal cardiomyocytes and postnatal cardiomyocytes quickly establish a nuclear MTOC and cell cycle arrest [17,32], we over-expressed Pcnt S in proliferating C2C12 myoblasts as they establish a nuclear MTOC during differentiation like cardiomyocytes [33]. In addition, it has been shown that Pcnt S is upregulated in adult skeletal muscle [15,16]. C2C12 cells were transfected with plasmids encoding Pcnt S-T2A-eGFP, Pcnt B-T2A-eGFP, or eGFP and cell cycle activity was analyzed based on nucleotide analog incorporation (5-ethynyl-2’-deoxyuridin (EdU)). The analysis of over 27.000 GFP-positive myoblasts from three independent experiments revealed that 33.2% control-transfected myoblasts incorporated EdU (Figure 5A-C). In contrast, only 20.8% Pcnt S-T2A-eGFP-transfected myoblast were EdU-positive (Figure 4B,C, p = 0.001). Notably, ectopic expression of Pcnt B-T2A-eGFP had no effect on EdU incorporation (33.9%, p > 0.05, Figure 5B,C). As an additional control, EdU incorporation was assessed in all experiments in the GFP-negative cells. The analysis of 218.819 GFP-negative cells revealed no difference in the percentage of GFP-negative/EdU-positive cells among the three different groups (control: 36.3%; Pcnt S-T2A-eGFP: 37.1%; Pcnt B-T2A-eGFP: 35.6%, all p > 0.05, Figure 5D,E).

**Figure 5.**
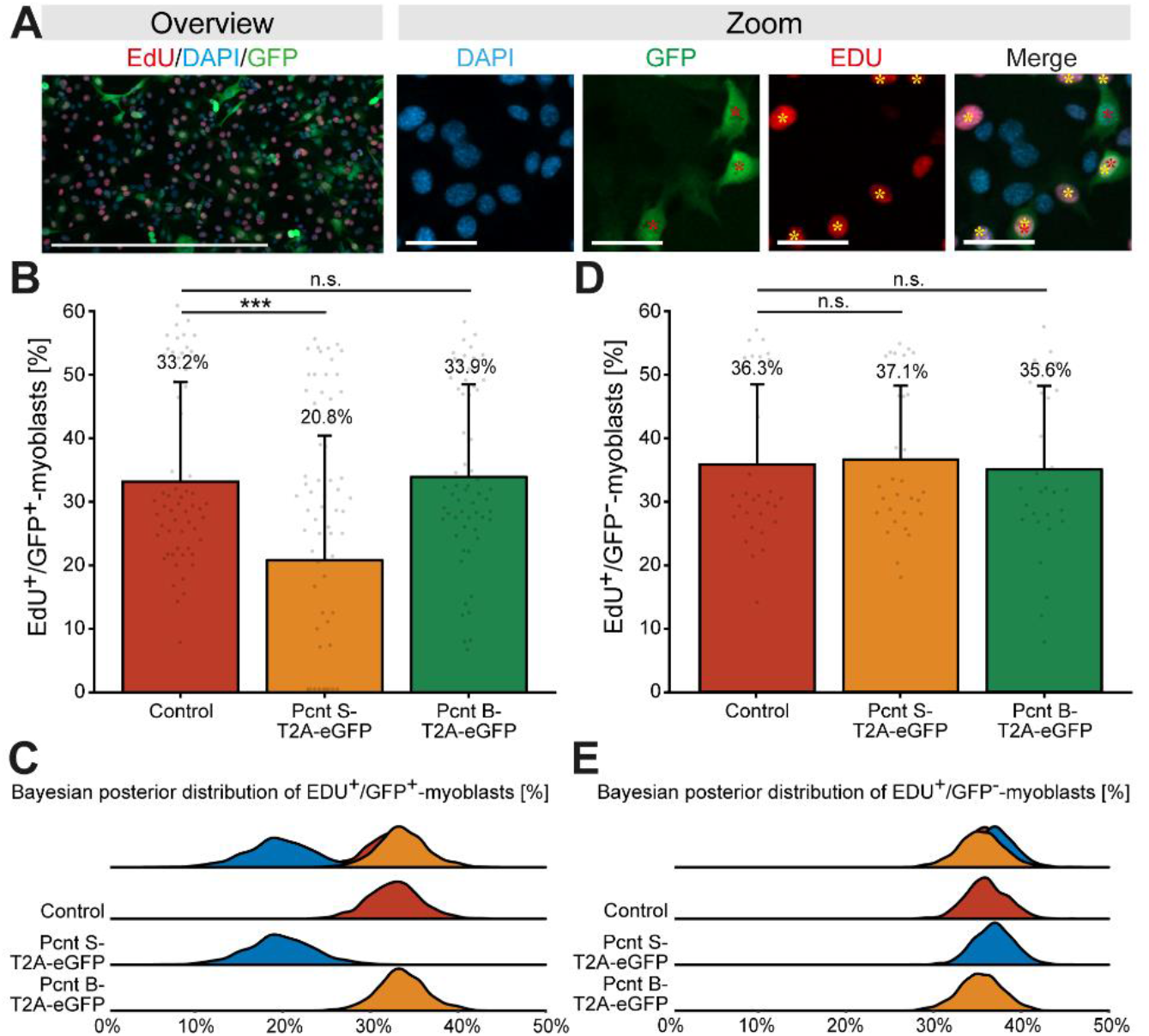
Ectopic expression of Pcnt S impairs DNA synthesis in C2C12 myoblasts. (**A**) Representative examples of transfected C2C12 myoblast (GFP, green) labeled with the nucleotide analog EdU (red). Nuclei were visualized with DAPI (DNA). (**B**,**D**) Quantitative analysis of EdU^+^/GFP^+^(B) and EdU^+^/GFP^−^ (D) myoblast upon control and Pcnt isoform expression as indicated. (**C**,**E**) Bayesian posterior distribution for the data in **B** and **D**. Red asterisk: GFP^+^ myoblast. Yellow asterisk: EDU^+^ myoblast. Scale bars: 50 μm (zoom) and 500 μm (overview). For the experiments ≥ 274 myoblasts were analyzed per experimental condition. Data are mean +- SD, n = 3, n.s.: p > 0.05, ***:p < 0.001.

Taken together, these results suggest that Pcnt S and thus alternative splicing of Pcnt might contribute to the establishment of cell cycle arrest in cardiomyocytes.

### 3.5. Pcnt S depletion enhances serum-induced cell cycle re-entry in cardiomyocytes

In order to determine whether depletion of Pcnt S increases the potential of cardiomyocytes to proliferate, P3 cardiomyocytes were stimulated with 10% FBS upon Pcnt S or Pcnt B+S depletion and progression into different cell cycle phases was assessed. Analysis of Ki67, a widely used marker to estimate the number of proliferating cells [34], revealed that siRNA-mediated depletion of Pcnt S significantly increased the number of Ki67-positive cardiomyocytes (control: 16.2%, 50 nM siPcntS: 26.4%, p < 0.001, 100 nM siPcntS: 28.2%, p < 0.001, 200 nM siPcntB+S: 26.2%, p < 0.001, Figure 6A-C). We have analyzed 28.496 cardiomyocytes from three independent experiments.

**Figure 6.**
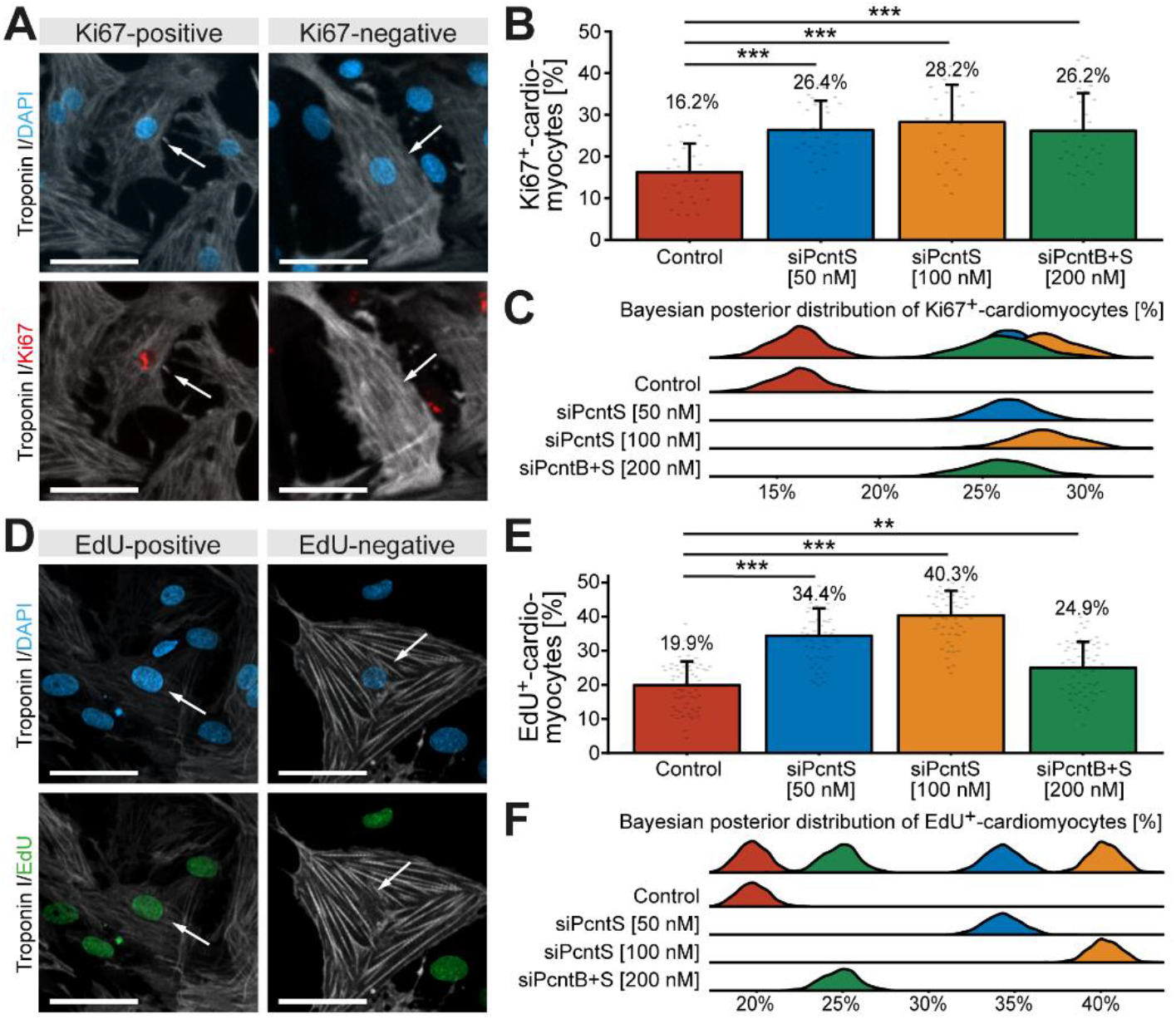
Depletion of Pcnt S increases cell-cycle activity and DNA synthesis in postnatal cardiomyocytes. (**A**,**D**) Representative examples of P3 cardiomyocytes (troponin I) stained for Ki67 (**A**) and incorporated EdU (**D**). Nuclei were visualized with DAPI (DNA). (**B**,**E**) Quantitative analysis of Ki67^+^ (**B**) and EdU^+^ (**E**) cardiomyocytes upon control and Pcnt isoform depletion as indicated. (**C**,**F**) Bayesian posterior distribution for the data in b and e. Scale bars: 50 μm. White arrows: cardiomyocyte nuclei. Data are mean ± SD, **: p < 0.01, ***: p < 0.001. For the experiments ≥ 15000 cardiomyocytes were analyzed per experimental condition.

As it has previously been indicated that Ki67 is not a very reliable marker to determine cardiomyocyte proliferation [11,35], we determined next the effect of Pcnt S depletion on 10% FBS-induced S phase entry by assessing EdU incorporation. The analysis of 56081 cardiomyocytes from three independent experiments showed that siRNA-mediated depletion of Pcnt S significantly increased the number of EdU-positive cardiomyocytes (troponin I-positive) (control: 19.9%, 50 nM siPcntS: 34.4%, p < 0.001, 100 nM siPcntS: 40.3%, p < 0.001, 200 nM siPcntB+S: 24.9%, p < 0.01, Figure 6D-F). Taken together, these data indicate that Pcnt S depletion enhances serum-induced cell cycle re-entry in cardiomyocytes into S phase.

### 3.6. Pcnt S depletion enhances serum-induced cell division in cardiomyocytes

To determine if Pcnt S depletion promotes also karyokinesis, we trained object classifiers (Methods, Section 2.8) to identify mitotic cardiomyocytes based on DNA staining patterns (DAPI, Figure S3A). We identified in all treatments together a total of 202 mitotic cardiomyocytes amongst 75387 analyzed cardiomyocytes from three independent experiments. While in the control group 0.21% cardiomyocytes entered mitosis (37/17479), depletion of Pcnt S resulted in 0.27% (52/18912, 50 nM siPcntS) and 0.31% (72/22973, 100 nM siPcntS) mitotic cardiomyocytes (Figure 7A).

**Figure 7.**
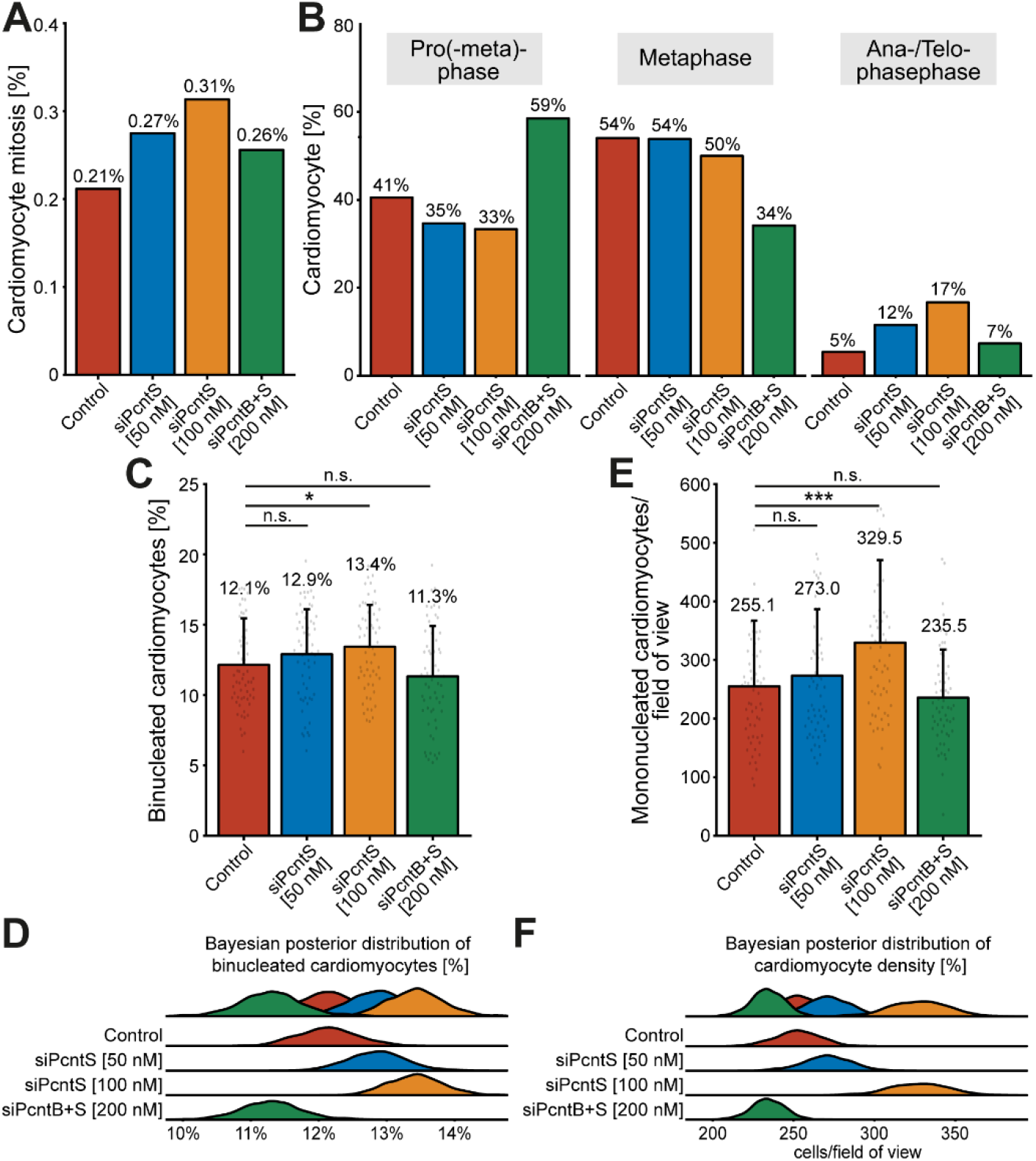
Pcnt S depletion enhances serum-induced cell division in cardiomyocytes. (**A**) Quantitative analysis of all mitotic cardiomyocyte within 3 independent experiments upon control and Pcnt isoform depletion as indicated. For the experiments ≥ 18.000 cardiomyocytes were analyzed per experimental condition. An object classifier was utilized that identifies mitotic cells based on DNA staining patterns. (**B**) Classification of the observed mitotic cardiomyocytes into pro(-meta)phase, metaphase, and ana-/telophase. (**C**,**E**) Quantification of binucleation (**C**) and mononucleated cardiomyocyte density (**E**). For the experiments ≥ 16023 cardiomyocytes were analyzed per experimental condition. Data are mean ± SD, n = 3, n.s.: p > 0.05, *: p < 0.05, ***: p < 0.001. (**D**,**F**) Bayesian posterior distribution for the data in c and e.

Depletion of Pcnt B+S resulted in to 0.26% (41/16023) mitotic cardiomyocytes (Figure 7A). Notably, a closer look at the percentage of mitotic cardiomyocytes in pro(-meta)-phase, metaphase and ana-/telophase suggested that Pcnt S-depleted cardiomyocytes have a higher potential to enter ana-/telophase and thus complete mitosis (control: 5% (2/37), 50 nM siPcntS: 12% (6/52), 100 nM siPcntS: 17% (12/72), 200 nM siPcntS: 7% (3/41), Figure 7B). However, even though the increase in ana-/telophase cardiomyocytes upon 10% FBS stimulation is more than 3-fold higher after Pcnt S depletion than in the control, the overall number of mitosis is very low. This might be due to the transient nature of mitosis and its very short duration (around 60 min) [10]. To further substantiate that Pcnt S depletion promotes 10% FBS-induced cell division, we determined whether Pcnt S depletion resulted in an increase in binucleated cardiomyocytes (Figure 7C,D and Figure S3B) or an increase in the number of mononucleated cardiomyocytes (Figure 7E,F). The analyses of 75387 cardiomyocytes from three independent experiments revealed that Pcnt S depletion has only a minor effect on serum-induced binucleation (control: 12.1%; 50 nM siPcntS: 12.9%, n.s.: p > 0.05; 100 nM siPcntS: 13.4%, p < 0.05; 200 nM siPcntB+S: 11.3%, n.s.: p > 0.05; Figure 7C,D). In contrast, Pcnt S depletion had a marked effect on cardiomyocyte number. The mean number of cardiomyocytes per field of view (FOV) after stimulation with 10% FBS was 255.1 in the control (Figure 7E,F). Pcnt S depletion resulted in an increase of ~7 and ~30% to 273.0 (50 nM siPcnt S) and 329.5 (100 nM siPcnt S, p < 0.001) cardiomyocytes per FOV, respectively (Figure 7E,F). Depletion of Pcnt B+S had no significant effect. Collectively, our data suggest that Pcnt S depletion enhances serum-induced mitosis in cardiomyocytes resulting in cell division.

## 4. Discussion

We conclude that expression of Pcnt S contributes to the cell cycle arrest in postnatal mammalian ventricular cardiomyocytes and Pcnt S depletion promotes cardiomyocyte proliferation. Several lines of evidence support these conclusions. First, ectopic expression of Pcnt S increases centriole splitting in ARPE-19 cells and inhibits DNA synthesis in C2C12 myoblasts. Second, the depletion of Pcnt S enhances serum-induced cell cycle activity, facilitates DNA synthesis, and promotes mitotic entry as well as anaphase transition in cardiomyocytes and results in a higher density of mononucleated cardiomyocytes.

Pcnt B is an anchoring protein of the MTOC that is located via its PACT domain at the centrosome [30,31]. In contrast, Pcnt S is found predominantly at the nuclear envelope [17]. The difference in localization is surprising, as both isoforms contain the C-terminal PACT domain [36]. Why Pcnt S is found predominantly at the nuclear envelope is unknown. However, it has been shown that “the partially translated PCNT nascent polypeptide starts to interact with the dynein motor complex once the dynein light intermediate chain 1 (LIC1)-interacting domain in the N-terminal half of PCNT is synthesized and folded” [37]. Pcnt S lacks this N-terminal LIC1 domain. Consequently, Pcnt B is the predominant Pcnt isoform at the centrosome due to an active transport mechanism enriching Pcnt B at the centrosome. This would explain our finding that both isoforms can bind to both centrosome and nuclear envelope (via their PACT domain) with different efficiencies due to an active transport of Pcnt B to the centrosome. As the primary structure and protein domains of Pcnt S are shared with the C-terminal part of Pcnt B, a transport mechanism to specifically enhance Pcnt S at the nuclear envelope is unlikely (Figure S1). Yet, it might be that the different 5’ UTR is utilized to transport the mRNA to the nuclear envelope followed by local translation.

The cell cycle arrest in cardiomyocytes might be a requirement for Pcnt S expression or might be the result of Pcnt S expression. Based on our data that ectopic Pcnt S expression induces centriole splitting and impairs cell cycle progression whereas Pcnt S depletion enhances serum-induced cell cycle progression, it appears that the cell cycle arrest in cardiomyocytes is a consequence of Pcnt S expression.

Yet, how Pcnt S affects cell cycle progression is unclear. Loss of centriole cohesion has been proposed to promote cardiomyocyte cell cycle exit and is associated with the upregulation of Pcnt S [17]. Importantly, ectopic expression of Pcnt S resulted in ARPE-19 cells in centriole splitting further indicating that Pcnt S actively induces cell cycle arrest in cardiomyocytes by inactivating the centrosomal MTOC. Finally, our data indicate that depletion of Pcnt S enhances the effect of factors stimulating cardiomyocyte proliferation. The depletion of Pcnt B, additionally to Pcnt S, seems to limit this stimulating effect through the induction of centriole splitting and subsequent cell-cycle inhibition. It is well known that serum stimulation induces in postnatal cardiomyocytes binucleation instead of cell division. Yet, here we show that serum stimulation upon depletion of Pcnt S resulted in increased cardiomyocyte cell division. Thus, it will be of interest to test in the future whether Pcnt S depletion can improve current strategies to regenerate the heart.

Pcnt S is upregulated during the time in which cardiomyocytes reorganize their cytoskeleton, potentially to meet increased functional demands. Thus, increased expression of Pcnt S might be a requirement for improved muscle function or a consequence of enhanced muscle function. While there is no obvious reason why Pcnt S should have a different function as Pcnt B, there are arguments for the need of a nuclear envelope MTOC which is facilitated by Pcnt S expression. An increasing work load of the heart during development requires an increasing contractile force generation by cardiomyocytes which causes more force on the nucleus. Thus, a nuclear envelope MTOC might be required to generate a microtubule-based cage around the nucleus to protect it from intracellular shear and compression forces to prevent DNA damage and maintain genome organization for a proper transcriptional program. On the other hand, the postnatal heart needs to respond to environmental changes (e.g. increased work load upon exercise) and this might require the rearrangement of the cytoskeleton. Similar is true for skeletal muscle. In addition, cardiomyocytes increase significantly in size during postnatal development and the formation of microtubules from a point source to a larger central radial source might be required to ensure the highly intracellular organization (e.g. sarcomeres, mitochondria, and t-tubules, invaginations of the muscle cell membrane) as well as efficient microtubule-based intracellular transport [38]. Notable, it has recently been demonstrated that the nuclear MTOC is required for cardiomyocyte hypertrophy [19]. In the future it will be important to determine whether Pcnt S has any specific role at the nuclear MTOC and to establish systems in which the nuclear envelope MTOC can be modulated and cellular organization and function can be assessed.

Taken together, this study suggests that expression of Pcnt S contributes to the cell cycle arrest in postnatal cardiomyocytes which, in turn, promotes a post-mitotic state. Given the increasing interest in cardiac regeneration and the role of non-centrosomal MTOCs in cell differentiation and function, understanding the establishment and function of non-centrosomal MTOCs and cell cycle arrest during development may reveal new mechanisms to regulate cell proliferation and function with implications not only for the treatment of striated muscle disease but in general for regeneration and cancer.

## Supplementary Materials

The following are available online after the main manuscript below: Figure S1: Overview of alternative splicing of PCNT and challenges for specific depletion of Pcnt S, Figure S2: Verification of Pcnt expression after ectopic expression in ARPE-19 adult retinal pigment epithelial cells, Figure S3: Representative examples of different cell cycle stages of cardiomyocytes.

## Author Contributions

Conceptualization, F.B.E., S.V., J.S.; formal analysis, J.S.; investigation, J.S., R.B., S.V.; writing—original draft preparation, J.S.; writing—review and editing, R.B., S.V., F.B.E.; visualization, J.S. and F.B.E.; supervision, S.V. and F.B.E.; project administration, F.B.E.; funding acquisition, J.S., S.V., F.B.E.. All authors have read and agreed to the published version of the manuscript.

## Institutional Review Board Statement

The investigation conforms to the guidelines from Directive 2010/63/EU of the European Parliament on the protection of animals used for scientific purposes. Extraction of organs and preparation of primary cell cultures were approved by the local Animal Ethics Committee in accordance to governmental and international guidelines on animal experimentation (protocol TS-9/2016 Nephropatho).

## Informed Consent Statement

Not applicable.

## Acknowledgments

The authors would like to thank the members of the working group of Prof. Felix Engel for support and discussion, Jana Petzold and Jennifer Redlingshöfer for technical support, and Kunsoo Rhee for the p3xFLAG-CMV10-eGFP-hPCNTB plasmid. The present work was performed in fulfillment of the requirements for obtaining the degree „Dr. med.” at the Friedrich-Alexander-Universität Erlangen-Nürnberg (FAU).

## Supplementary Material

**Figure S1.**
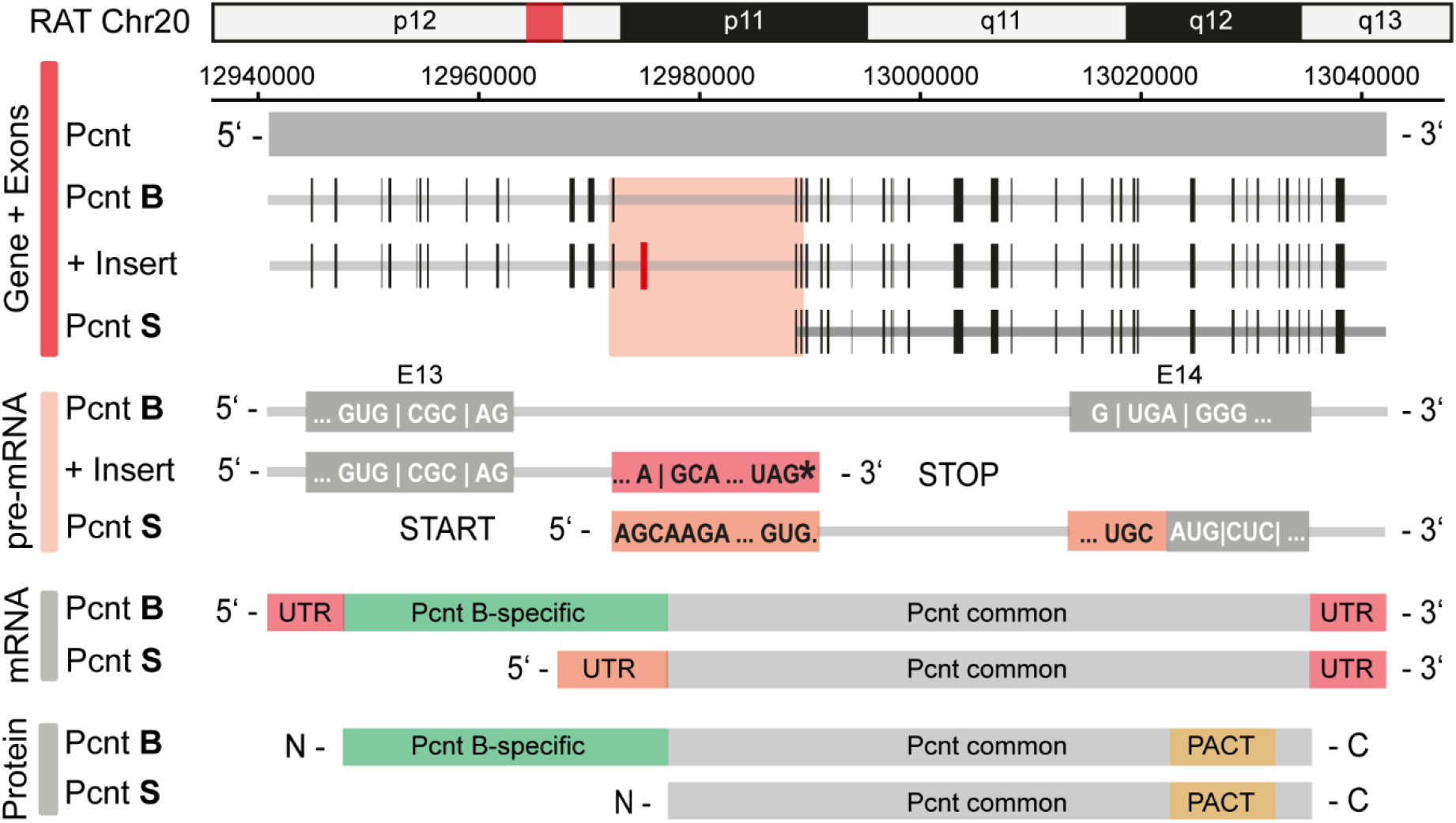
Overview of alternative splicing of PCNT and challenges for specific depletion of Pcnt S. Alternative splicing of rat Pcnt results in the insertion of an intronic region in the Pcnt gene between exon 13 and 14, which becomes the first exon encoding the isoform Pcnt S (Gene + Exon). This exon contains a stop codon and consequently initiation of Pcnt S translation occurs in a downstream start codon resulting in the truncation of the first 232 amino acids at the N-terminal region of the longer Pcnt B isoform. The new 5’-UTR is constituted of the sequence between stop codon and new start codon, in part consisting of a coding sequence for Pcnt B (pre-mRNA). The lack of specific regions in the expression of Pcnt S presents a major challenge for specific Pcnt S depletion and precludes Pcnt S-specific antibody staining. To specifically deplete Pcnt S we used a siRNA against the specific part of the 5’-UTR of the Pcnt S mRNA (mRNA).

**Figure S2.**
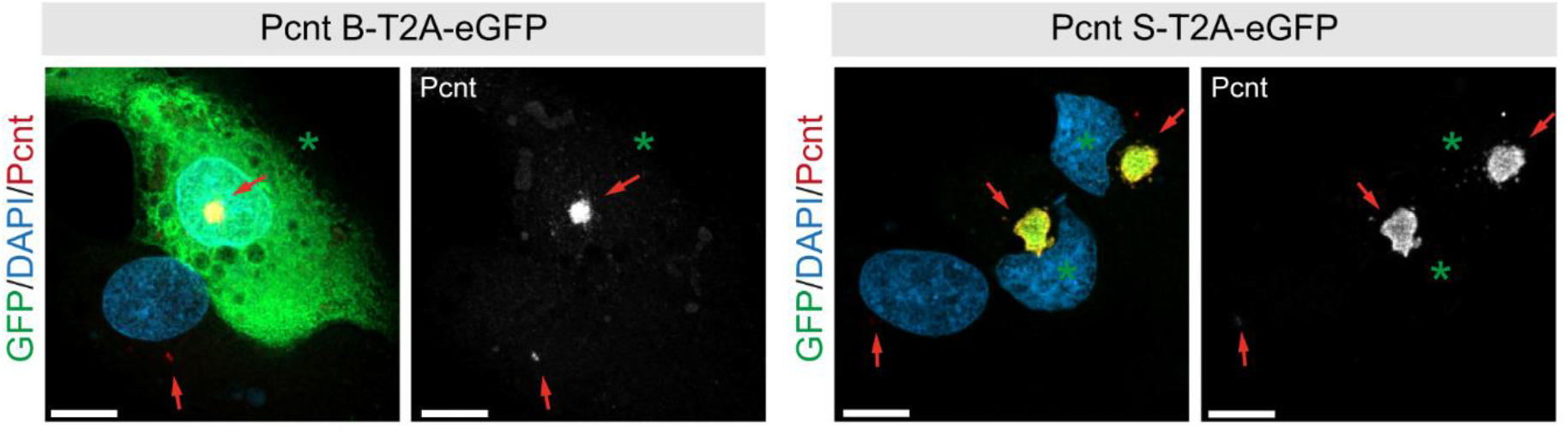
Verification of Pcnt expression after ectopic expression in ARPE-19 adult retinal pigment epithelial cells. Representative examples of ARPE-19 cells transfected with bi-cistronic plasmids encoding Pcnt B-T2A-eGFP or Pcnt S-T2A-eGFP (GFP, green) and immunostained with anti-pcnt antibody (red). Nuclei were visualized with DAPI (DNA, blue). Green asterisk: GFP+ cell, Red arrow: Pcnt staining. Scale bars: 10 μm.

**Figure S3.**
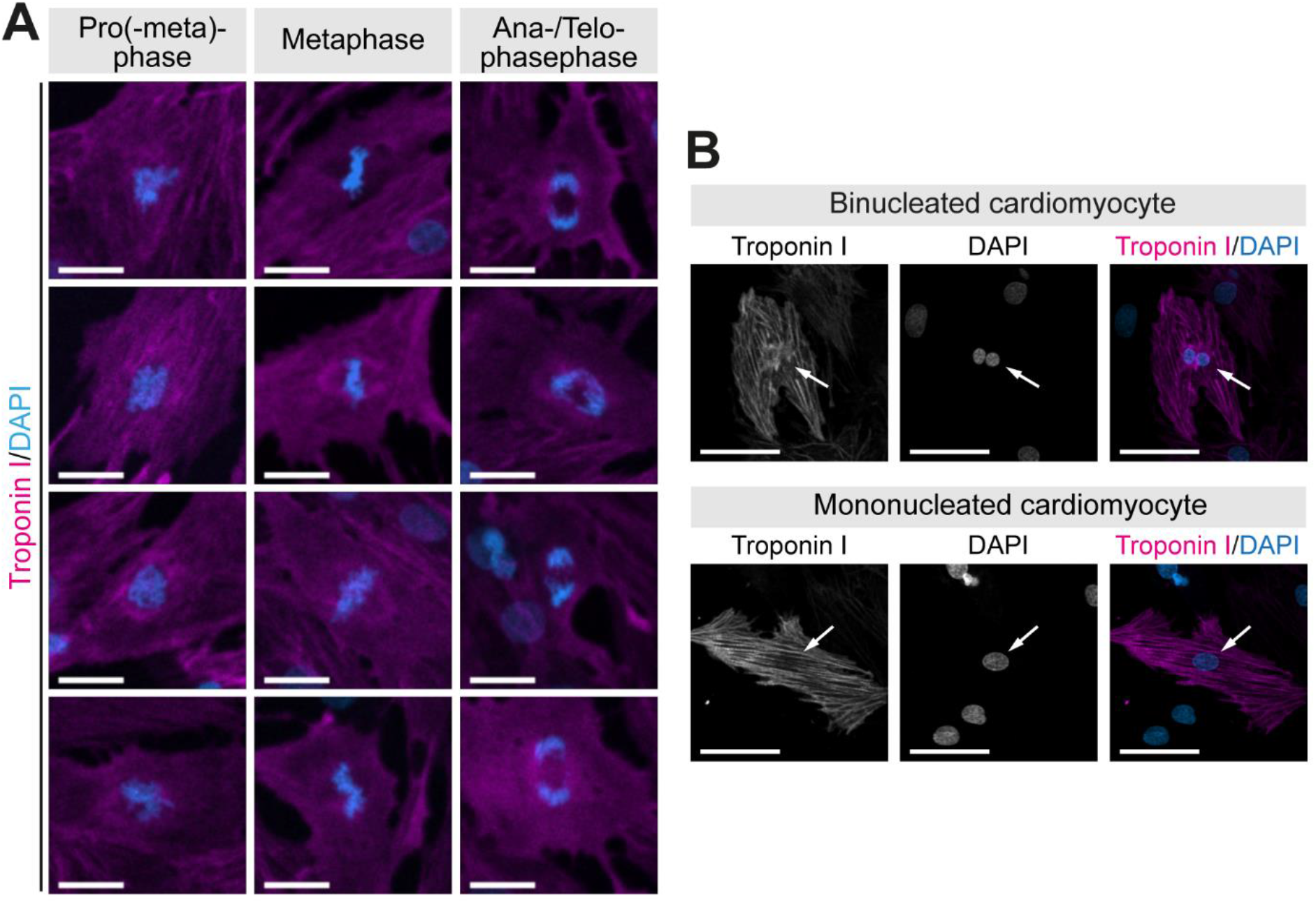
Representative examples of different cell cycle stages of cardiomyocytes. (**A**) Representative examples of cardiomyocytes (troponin I) in pro(-meta)phase, metaphase and ana-/telophase. Chromosomes were visualized with DAPI (DNA). Scale bars: 20 μm. (**B**) Representative examples of mono- and binucleated cardiomyocytes (troponin I). Nuclei were visualized with DAPI (DNA). White arrows: cardiomyocyte nuclei. Scale bars: 50 μm.

